# Global 5-methylcytosine-RNA disruption reduces the vectorial competence to DENV2 of heatwave-exposed *Aedes aegypti* mosquitoes

**DOI:** 10.1101/2024.03.14.585075

**Authors:** Fabiola Claudio-Piedras, Benito Recio-Tótoro, Humberto Lanz-Mendoza, Jorge Cime-Castillo

**Author notes:** **Correspondence:** Jorge Cime Castillo.

## Abstract

Heatwaves are an increasingly common environmental event linked with climate change. Abnormally high heatwave temperatures can affect several mosquito vector traits that are determinants of pathogen transmission. Understanding how these mosquitoes adapt to high heat is vital for global public health. RNA methylation, a key cellular mechanism in stress response and adaptation, remains understudied in mosquito vector competence and heat stress responses. This study investigates the role of RNA methylation in mosquito responses to heatwaves and its influence on DENV2 vector competence. Heatwave-exposed and DENV-infected mosquitoes presented lower survivorship and lower antiviral transcriptional response, developed high infection rates, and increased the life expectancy of infected mosquitoes during the period of highest virus transmissibility. In contrast, inhibition of RNA methylation in heatwave-treated mosquitoes increased survivorship and the antiviral transcriptional response, reducing infection prevalence from 78% to 37%. These results indicate that the RNA methylation background in mosquitoes favors vector competence for DENV2 during a heatwave exposure, and points towards possible interventions to countermeasure the effect of climate change on DENV transmission.

## 1 Introduction

Extreme weather events associated with climate change, such as heatwaves, are becoming more frequent and expose disease vector mosquitoes to changes in temperature, humidity, and food availability (1,2). As ectotherm animals, mosquitoes’ physiological and ecological traits are heavily dependent on temperature (3,4). Temperatures exceeding the thermal optima for the mosquito’s biological cycle represent sources of stress that lead to physiological changes ranging from increased metabolic activity to pathological states and insect mortality (5–7). hese physiological changes are transient and enable the functionality of essential cellular components during the stress period and the resumption of normal cellular activities during the recovery period (4,8). In *Aedes aegypti*, an increase in average temperature enhances vector survival, accelerates the extrinsic incubation period for dengue virus, promotes biting behavior, and increases insecticide resistance (9–11).

The remarkable plasticity of mosquitoes in the face of environmental adversity is evidenced by their persistence in a changing climate, ample global distribution, and extensive species radiation (2). RNA methylation is one of the fundamental molecular mechanisms contributing to cellular adaptation to environmental changes and stressful stimuli (12–17). However, little is known about the contributions of this mechanism to mosquito vector competence and response to thermal stress (18–23). Cytosine methylation in RNA is a post-transcriptional regulatory mechanism conserved in archaea, prokaryotes, and eukaryotes, affecting nearly all aspects of RNA processing (Reviewed in (24). The chemical modification of cytosine at the C5 position is catalyzed by methyltransferases, which transfer a methyl group from the donor S-adenosylmethionine (SAM) to the C5 position of cytosine, converting it to 5-methylcytosine (5mC). DNMT2 is the most conserved methyltransferase among eukaryotic organisms and has catalytic activity on both DNA and RNA (24,25). Nucleoside analogues, such as azacytidine inhibit DNMT2 activity. Substitution of C5 in cytosine for N5 in azacytidine inhibits C5 methylation and disrupts beta-elimination, leaving the enzyme covalently bound to the nucleic acid and subsequently degraded (Reviewed in: (26–28). 5mC is removed by demethylases and recognized by methyl group-binding proteins, which exert regulatory functions after binding (24).

In dipterans, the biological processes involving DNMT2 include: 1) the maintenance of the lifespan of adult flies (29); 2) positive regulation of the antiviral defense response (30); 3) positive regulation of the innate immune response (31,32); 4) improved tRNA stability (33,34); 5) response to heat shock-induced stress (23,33,35); 6) response to oxidative stress (35). The content of 5mC varies with physiological changes during development, metamorphosis, and aging, as well as changes in methylation profiles induced by nutritional deficiencies, oxidative stress, thermal stress, and infection (36).

*A. aegypti* is the primary mosquito vector of dengue virus (DENV) worldwide (37). DENV transmission begins when a female mosquito feeds on viremic blood from a mammalian host, acquiring infectious viral particles along with the blood. DENV efficiently enters the epithelial cells of the mosquito’s gut and establishes an infection that spreads within 1 to 2 days throughout the mosquito’s gut (38–40). In the competent mosquitoes’ midgut, active viral replication occurs, which is crucial for subsequent dissemination and transmission of the virus. Starting from day 4, viruses released from the gut must traverse several tissues to reach the salivary glands, where they are secreted in the saliva around day ten and can be transmitted to humans through subsequent bites (38–41). In this study, we used the *A. aegypti*-DENV2 model to investigate the role of RNA methylation in response to thermal stress, viral infection, and vector competence.

## 2 Materials and methods

### 2.1 Mosquitoes

*Aedes aegypti*, Rockefeller strain mosquitoes were handled under the Instituto Nacional de Salud Pública (INSP) biosafety and ethics guidelines. The mosquitoes were reared and kept at standard conditions of 28°C, 60-80% RH, and a 12/12 h light-dark cycle. Recently hatched larvae were reared in 693 cm^2^ plastic containers at a density of 200 larvae per 2.5 L of tap water previously left overnight to reach the insectary temperature. Larvae were fed a standard mix of yeast extract, lactalbumin, and grounded mouse pellets (1:1:1) in dechlorinated water until pupation. Pupae were collected in 6 cm in diameter plastic cups at a density of 500 pupae per cup in ∼50 ml of tap water and let to emerge in 4 L adult plastic containers. Adult mosquitoes were kept under 10% sugar solution-dampened cotton pads replaced every 24 h. All experimental mosquitoes received antibiotic treatment (PSN: 5000 U/ml of penicillin, 5 mg/ml of streptomycin, and 10 mg/ml of neomycin) added to the sugar solution from the fifth day post-emergence (dpe) until the end of experimentation.

### 2.2 Virus

Dengue virus serotype 2 (DENV2) strain New Guinea was propagated by alternated passages between C6/36 *Aedes albopictus* and BHK 21 hamster cell lines before infecting mosquitoes. Briefly, DENV2 was activated by inoculating monolayer cultures of C6/36 cells at 80% confluence for 2 h at 28°C. The inoculum was then removed and the cells maintained in Schneider medium with 2% of heat-inactivated fetal bovine serum (hiFBS) for 5 to 7 days at 28°C. The supernatant containing the virus was collected, centrifuged, and filtered through a 0.2 µm membrane for its inoculation in BHK 21 cells at 80% confluence for 2 h at 37°C and 5% CO_2_. After removal of the initial inoculum, BHK 21 cells were kept at 37°C and 5% CO_2_ in DMEM medium with 2% hiFBS. The supernatant containing virus was collected after 5 to 7 days for its centrifugation and filtration as stated above. This procedure was repeated three times and the virus collected from the last BHK 21 cell passage was used for infecting mosquitoes.

### 2.3 Mosquito infections

The standard membrane feeding assay was used to infect seven days post-emergence (dpe) mosquitoes previously starved for 12 h and treated with PSN from the previous 48 h. Infections were carried out with a mixture of 10^5^ copies of DENV2 viral RNA per ml of supernatant and rabbit blood (1:1, v/v) at 37°C for 30 min in the dark. Mock infections were used as control by replacing the DENV2-containing supernatant with non-inoculated BHK 2-cell supernatant. Unfed and partially engorged females were excluded from the experiments. Immediately after blood-feeding, the mosquitoes were transferred to incubators set to either 28°C for the control groups or 34°C for the heatwave groups. Wet filter papers in plastic cups were placed 48 h after blood-feeding to allow for oviposition; collecting the eggs 72 h after.

### 2.4 RNA methylation inhibition

The 5-methylcytosine (5mC) inhibitor azacytidine was given to the mosquitoes during the blood-feeding by adding it to the DENV2/blood mixture at a final concentration of 50 µM. The azacytidine treatment continued for five days but now added to the 10% sugar and PSN solution-dampened cotton pads. The treatment was administered every 12 h on new cotton pads.

### 2.5 Heatwave

The mosquitoes were exposed for five days to a heatwave right after blood-feeding in an incubator (min = 33.6°C, max = 38.2°C, Supplementary Figure 1A), 60 to 80% RH and a 12/12 h day/night cycle. The temperature and relative humidity were recorded every 3 min. Cotton pads with 10% sugar-PSN solution were replaced every 12 h during the heatwave. After the simulated five-day long heatwave, the mosquitoes were transferred to 28°C, 60 to 80% RH, and 12/12 h day-night cycle. The cotton pads dampened with 10% sugar-PSN solution were then replaced every 24 h.

### 2.6 Mosquito survival

After blood-feeding, daily survival curves of 100 mosquitoes from each experimental condition (DENV2- or mock-infected, azacytidine-treated, and heatwave-exposed mosquitoes) were recorded. The survival curves were evaluated for scale and shape effects. The scale effects were determined by calculating the characteristic life (α) as the age at which ≈37% of the population is still alive (*s(α)=exp*(*-1*)), and the median survival, defined as the age at which 50% of the population is still alive. Shape effects were determined by rescaling the age (x) by (α) for each survival curve (42). Life expectancy was calculated using the following formula:

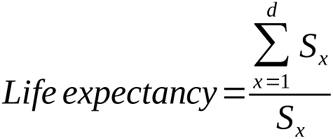

Where *x* represents the day of interest, *d* the day at which the last mosquito of the cohort died, and *S* the proportion of surviving mosquitoes at day *x*.

### 2.7 RT-qPCR

The transcriptional response of *dnmt2*, *hsp70*, *r2d2*, *dicer1*, *dicer2*, and *ago1* to DENV2, exposure to a heatwave, and the azacytidine treatment, was evaluated by qPCR in midgut (0, 1, and 4 days post-infection, dpi), head, thorax and abdomen (7 dpi) samples coming from 90 mosquitoes per group. For this, midguts were dissected and washed several times to remove blood remains and the head, thorax, and abdomen separated while still ice-cold. RNA extractions were carried out by the TRIzol (Thermo Fisher Scientific) protocol. RNA samples were stored at -20°C until further use.

For cDNA synthesis, RNA extracts (1 μg) were treated with DNase I and 500 ng/μl of OligodT were added. After 10 min at 70 °C, the master mix (1mM dNTP’s, 0.01mM DTT, 20 units of RNase inhibitor and reverse transcriptase buffer) and 200 U of M-MLVRT were added.

*dnmt2*, *hsp70*, *r2d2*, *dicer1*, *dicer2*, *ago1*, and *RNAsaP* genes were amplified by mixing 10 ng of cDNA, 0.5 μM oligonucleotides (Supplementary Table), and Master Mix SYBR Green. Amplifications were carried out on a RotorGene (Qiagen) thermalcycler following the next program: 95°C / 2 min; (95°C / 2 s; 60°C / 20 s) 40X. The amplification efficiency was determined with LinRegPCR. Data were normalized to *RNAsaP* gene amplification.

### 2.8 Viral RNA detection in mosquito samples

Viral RNA was quantified in single mosquitoes (n=30) at 7 dpi. RNA was extracted by the TRIzol protocol and stored at -20°C until further use. For cDNA synthesis, RNA extracts (500 ng) were treated with DNase I and 500 ng/μl of random hexamers were added. After 10 min at 70°C, the master mix (1mM dNTP’s, 0.01mM DTT, 20 units of RNase inhibitor and reverse transcriptase buffer) and 200 U of M-MLVRT were added. Quantification of viral RNA in mosquito samples was performed by qPCR, in relation to a synthetic DENV standard curve comprised of seven 10-fold serial dilutions from a stock of known concentration. Viral RNA was amplified on a RotorGene (Qiagen) thermalcycler following the next program: 95°C / 3 min; (95°C / 5 s; 60°C / 30 s) 40X. The amplification efficiency was determined with LinRegPCR.

### 2.9 HPLC-FLD

The content of 5mC in RNA was quantified by High Performance Liquid Chromatography coupled with a Fluorescent Light Detector following the methodology of Lopez-Torres, et al. (43). Ice-chilled head, thorax, abdomen, and midgut samples were prepared as described above making sure no blood remained in any of the samples. RNA from each body segment was extracted by the TRIzol protocol and stored at -20°C until further use. Total RNA was digested with RNase I and Nuclease P1 by mixing 2 µg of total RNA with hydrolysis buffer consisting of 20 mM CH_3_COOH, 20 mM glycine, 5 mM MgCl_2,_ 0.5 mM ClZn, 0.2 mM ClCa, 2 U of RNase I, and 0.2 U of Nucleasa P1 in a final volume of 47.5 µl. Reactions were incubated overnight at 37°C and heated at 95°C for 5 min followed by rapid cooling at 4°C. Then, 10 mM NaOH and 0.2 U of alkaline phosphatase were added for further incubation at 37°C for 2 h. Digested RNA samples were derivatized with 2-bromoacetophenone by first drying the samples in a SpeedVac SC100 (Savant) at 65°C for 25 min. Samples were then reconstituted in 135 µl of DMF with 3.7% of CH_3_COOH and 20 µl of 0.5 M 2-bromoacetophenone in DMF and 25µl of a saturated solution of Na_2_SO_4_ were added. Derivatization was left to occur at 80°C for 120 min covered from light.

The chromatography was carried out in an Agilent Series 1100 HPLC system with a Sorbax C18 (250 x 4.6 mm, 5 µm) column set to 30°C and a Supleco precolumn. Samples were diluted 1:1 with water and 20 µl injected into the system. Four mobile phases were used: water with 5% methanol (A), acetonitrile (B), 0.4% trifluoroacetic acid (C), and methanol (D). 5mC was separated from other nucleosides with the following gradient: starting from 75% A, 5% B, 0% C, 20% D; the gradient was modified from 0-7 min to 49% A, 10% B, 13% C, 28%D; from 7-11 min to 47% A, 12% B, 13% C, 28% D; from 11-14 min to 12% A, 15% B, 13% C, 60% D; and maintained from 14-16 min at 12% A, 15% B, 13% C, 60% D at a flow rate of 350 µl/min. Fluorimetric detection was achieved at 306/378 nm excitation/emission wavelengths, respectively.

### 2.10 Statistical Analysis

All variables (viral load, 5mC content, gene differential expression, survivorship, and life expectancy) from all experimental conditions and treatments (Mock-infected, DENV2-infected, heatwave-exposed, and azacytidine-treated mosquitoes) were analyzed by principal component analysis. The conditions and treatments were further clustered hierarchically to find patterns in the data. These analyses were conducted in R v 4.2.0 and plotted with the ggbiplot and ape packages. The following statistical analyses were conducted in Prism 6. Survival curves were analyzed with the Log-rank test. Viral RNA load, transcriptional response and 5mC content were analyzed by ANOVA followed by Tukey’s multiple comparison test. Correlation analysis of the transcript’s relative expression (log2-transformed 2^-Δct^ normalized with *RNAsaP* mRNA) were conducted by the Pearson correlation coefficient and plotted in Cytoscape with a cut-off threshold of *p* ≤ *0.05*.

## 3 Results

### 3.1 *dnmt2* is a stress-response element in *A. aegypti*

To explore whether RNA methylation is involved in the stress response, we first analyzed the expression of *dnmt2* in DENV2-infected, heatwave-exposed, and azacytidine-treated mosquito midguts. The mosquitoes were treated with azacytidine three days before being infected with DENV and subsequently exposed to heatwave (min = 33 °C, max = 38 °C) for 24 h, time at which the samples were taken. As seen in Figure 1A, all treatments and their combinations increased the expression of *dnmt2* in relation to untreated mosquito midguts. Notoriously, DENV2 infection in combination with the azacytidine treatment (DENV2+A), increased the expression of *dnmt2* more than 21 fold, while its counterpart DENV2-infected (DENV2) or azacytidine-treated only (A) mosquito midguts, upregulated *dnmt2* similarly to each other 13 and 11 fold in relation to control midguts, respectively. Samples midguts of mosquitoes heatwave-exposed (H and DENV2+H) upregulated *dnmt2* to a lesser extent between 5 and 7 fold compared to midguts of mosquitoes untreated. It is noteworthy that the effects of azacytidine and the heatwave on *dnmt2* expression when the mosquitoes are also infected with DENV2, are opposite; azacytidine upregulated *dnmt2* 8 fold relative to DENV2-infected only mosquito midguts (DENV2+A vs. D), whereas heatwave-exposed mosquitoes expressed *dnmt2* 8 fold lower (DENV2+H vs. D) and to levels similar to those of heatwave-exposed only mosquitoes (H).

**Figure 1.**
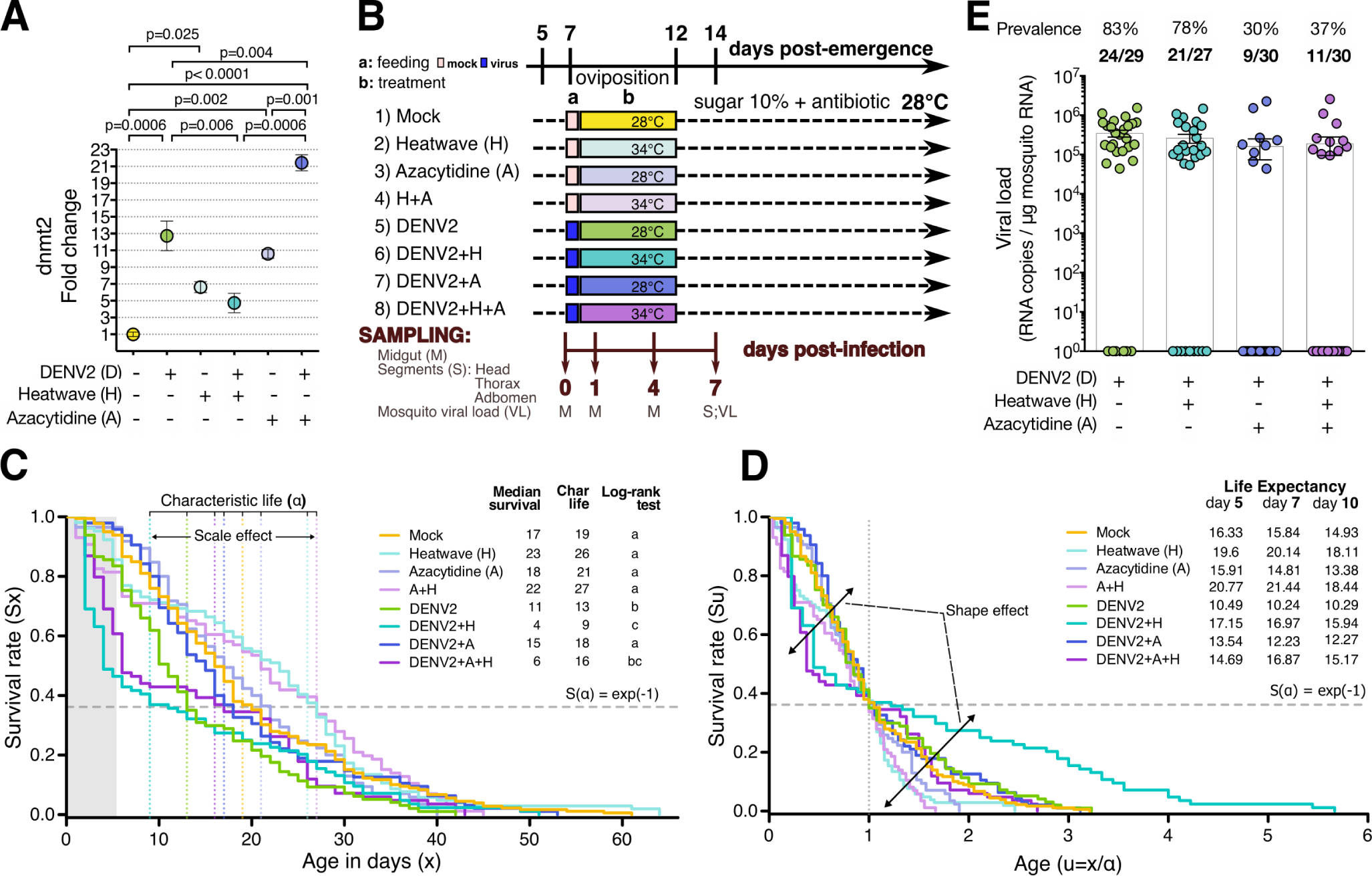
Effect of heatwave exposure and azacytidine treatment on survival and infection prevalence of *A. aegypti* mosquitoes challenged with DENV2. **(A)** Expression of *dnmt2* in mosquitoes exposed to a heatwave or treated with azacytidine. Relative expression of *dnmt2* in *A. aegypti* females 24 h following DENV2-infected blood feeding, exposure to a heatwave (H) or 50 µM azacytidine for 3 days, by qPCR. Normalization with *RNase P* and analysis by the 2-Δ^Ct^ method. Representation of the mean ± SEM of two independent experiments with three replicates each. **(B)** Experimental design. *A. aegypti* females were grouped based on exposure to dengue virus (DENV2), heatwave (H) or azacytidine treatment (A). Eight groups were formed considering all the treatment combinations. The 5 post-emergence (dpe) groups of females were treated daily with antibiotics in sucrose solution until the end of the experiment. 7 dpe mosquitoes were fed (a) with mock-infected blood (pink) or DENV2-infected blood (blue) and immediately treated (b) for 5 days with 50 µM azacytidine (A) or exposed to a heatwave (H). Survival, transcriptional response (*dnmt2*, *hsp70*, *r2d2*, *dicer1*, *dicer2* and *ago1*), 5mC content, infection prevalence and DENV2 viral load were analyzed. Samples were obtained from midgut, abdomen, thorax, head and whole mosquito. Samples were taken for total RNA extraction at 0, 1, 4 days post-feeding (dpf), at 7 dpf from abdomen, thorax and head, and from whole individual mosquitoes at 7 days post-infection (dpi). **(C)** Survival of mosquitoes exposed to DENV2, a heatwave or azacytidine treatment. Depiction of characteristic life span and median survival. Survival rates were determined in groups of 100 mosquitoes on two separate occasions and analyzed by the log-rank test; different letters denote statistical differences at p ≤ 0.05. Scale effects were determined by calculating the characteristic life (α) as the age at which ≈37% of the population is still alive (s(α)=exp(-1)), and the median survival, defined as the age at which 50% of the population is still alive. **(D)** Shape effects on survival curves of mosquitoes exposed to DENV2, a heatwave, or azacytidine treatment. Shape effects were isolated from scale effects by relativizing age (x) by characteristic life span (α). Life expectancy is equal to the sum of survival in days from the age of interest (*x*) to the day the last mosquito died (*d*), divided by the proportion of surviving mosquitoes at the day of interest (*x*). Life expectancy at days 5, 7, and 10 following feeding is shown. **(E)** Prevalence of infection and 7 dpi viral loads in mosquitoes exposed to heatwave or azacytidine treatment. Total RNA was extracted from approximately 30 individual mosquitoes per group to determine the viral load per mosquito by qPCR. The prevalence of infection was calculated as the percentage of infected mosquitoes. The ratio of infected mosquitoes/mosquitoes tested is shown.

These results gave us insights into the potential role of *dnmt2* in the response to a DENV2 infection and thermal stress, so we proceeded to explore the effects of cytosine methylation in RNA on the capacity of the mosquito to cope with a DENV infection in a scenario where a heatwave is also happening. For this, we devised the experimental design depicted in Figure 1B, wich consisted of eight experimental groups: 1) Mock-infected, 2) heatwave (H), 3) Azacytidine (A), 4) H+A, 5) DENV2, 6) DENV2+H, 7) DENV2+A, and 8) DENV2+H+A. All groups were treated daily with antibiotics starting at 5 dpe, and blood-fed with either DENV2-infected blood or mock-inoculated blood at 7 dpe. For its corresponding groups, 50 µM azacytidine was added to the blood for feeding and the treatment continued for five days by adding it to the sugar solution used as maintenance feeding. After a 30 min blood feeding period, the mosquitoes were transferred to incubators set at 28°C (min = 27.5°C, max = 28.5°), the optimal temperature for *A. aegypti*, or at 34°C (min = 33.6°C, max = 38.2°C) until 12 dpe to simulate a 5-day lasting heatwave. Mosquito midgut samples were taken at 0, 1, and 4 dpi, and head, thorax, and abdomen samples were taken at 7 dpi to evaluate the expression profile of *dnmt2*, heat-shock protein 70 (*hsp70*), the iRNA pathway, and the 5-methylcytosine content in RNA. Whole mosquito samples were taken at 7 dpi to determine the viral load. The rest of the mosquitoes were maintained until the last mosquito died to obtain survival curves for each experimental group.

### 3.2 Exposure to a heatwave increases the life expectancy of infected mosquitoes during the period of highest virus transmissibility

The mosquitoes infected with DENV2 exhibited a notably reduced survival rate, particularly within the first 12 dpi (Fig 1C). This decrease in survival was further exacerbated when the infected mosquitoes were exposed to a heatwave (DENV2+H, Log-rank test, *p*< 0.05). Interestingly, azacytidine recovered the survival rate of DENV2-infected mosquitoes (DENV2+A) to that of the mock-infected mosquitoes. A similar scenario happened with DENV2-infected and heatwave-exposed mosquitoes (DENV2+H+A), in which the azacytidine treatment increased the survival rates, albeit non-significantly. The tendency of heatwave-exposed mosquitoes (H and DENV2+H) to reach older ages is remarkable despite having a lower survival rate during heatwaves.

To explore this further, we analyzed the shape and scale effects of the experimental conditions on the survival rate of the mosquitoes. The scale effect refers to changes in the survival curves that result in increased or decreased lifespan. Whereas shape effects indicate changes in the survival rates at discrete ages, e.g. mosquitoes die younger or become older without necessarily changing their lifespan. Scale effects are usually measured as the characteristic life or the median survival, and shape effects are measured as the bendiness of the survival curves. Regarding the scale effects, the characteristic life (α), defined as the age (x) at which ≈ 63% of the population has died (Figure 1C), is considerably lower for DENV2-infected mosquitoes (DENV2), and even lower for those that were also exposed to a heatwave (DENV2+H). Mock-infected mosquitoes, DENV2-infected and azacytidine-treated mosquitoes (DENV2+A), as well as the DENV2-infected, azacytidine-treated and heatwave-exposed mosquitoes (DENV2+H+A) showed similar characteristic life. In the case of heatwave-exposed mosquitoes, treated or not with azacytidine (H, and H+A), the characteristic life is higher, indicating that heatwave-exposed mosquitoes live longer. However, the changes in shape need to be considered to get a clearer view of the effects of the treatments. So, to isolate the shape effects, the scale effect was filtered out by rescaling the age (x) by α (rescaled age = u = x / α, Figure 1D). There were considerable changes in shape earlier in life that correspond to the heatwave exposure (H, H+A, DENV2+H, and DENV2+H+A), indicating that the mosquitoes died younger, as one can expect. However, the DENV2-infected mosquitoes that survived exposure to a heatwave (DENV2+H), were able to reach older ages, as indicated by the increased survival rates later in life. Treatment with azacytidine reverses the extent of heatwave-induced survival in infected mosquitoes during the period of peak viral transmissibility, supporting that 5mC is involved in acclimatization. Although these mosquitoes at some point reached the survival rates of the uninfected mosquitoes (Figure 1C), the survival rates plumbed soon after at ≈ 25 days old, unlike the heatwave-exposed and infected mosquitoes (DENV2+H), which reached and maintained the survival rates to that of the non-infected mosquitoes. It is noteworthy that DENV2 mainly caused scale alterations in the survival rates (Figure 1C); whereas it only exacerbated the shape effects of the heatwave during the heatwave (DENV2+H and DENV2+H+A vs. H and H+A).

As mentioned earlier, azacytidine caused the infected mosquitoes (DENV2+A) to have characteristic life similar to the uninfected mosquitoes (Mock), even if exposed to a heatwave (DENV2+H+A). Azacytidine counteracts the scale effects introduced by DENV in combination with a heatwave (Figure 1C; DENV2+H+A). In terms of shape-related effects, azacytidine eliminated the shape alterations induced by the heat-wave later in life only in infected mosquitoes (DENV2+H vs. DENV2+H+A). This finding has important implications in DENV transmission dynamics because, in a scenario where a mosquito population experiences a heatwave, there is a higher chance for the surviving mosquitoes to reach older ages (shape effect) if infected, and live longer (scale effect) if not infected. Noteworthy is the effect of azacytidine, as it counteracts the shape deviations induced by a heatwave, especially when combined with DENV, nevertheless, it also increases the characteristic life, making the mosquitoes to live longer.

We calculated the life expectancy for each experimental group to determine how much longer the treated mosquitoes are expected to live. Since DENV takes between 7 and 14 days to reach the salivary glands, mosquitoes that are expected to live longer have a greater probability of transmitting the virus. Figure 1D depicts such live expectancy curves, the heatwave-exposed mosquitoes have a greater life expectancy, which, at 7 days and 10 days post-infection, are expected to live between 17 to 16 more days. In contrast, DENV-infected mosquitoes, at 7 and 10 dpi are expected to live for 10 more days. So, mosquitoes that have experienced a heatwave are expected to live 4 to 6 extra days, and since the gonotrophic cycle takes 3 days, these mosquitoes could potentially have the opportunity to bite at least one more host and lay another batch of eggs.

### 3.3 Azacytidine reduces the prevalence of DENV infection in *A. aegypti*

To evaluate the effect of azacytidine treatment and exposure to a heatwave, we individually tested 30 mosquitoes per condition at 7 dpi and quantified by RT-qPCR the viral load as the number of viral RNA copies per µg of total RNA. Azacytidine was the only experimental condition that impacted on the viral infection (Figure 1E). It is striking that the differences were observed in the infection prevalence, showing a 53% reduction in the number of infected mosquitoes, but not in the viral load of those still infected mosquitoes. This suggests of an all-or-none mechanism, in which the virus is either successfully fought back or allowed to thrive. Although most evident in the azacytidine-treated mosquitoes, this was also the case for all experimental conditions. These results suggest that RNA methylation favors DENV infection.

Determining the viral load and prevalence also allowed us to test if the changes in the survival rates were due to differences in the infection. We know that DENV causes scale effects on the mosquito survival rate, lowering it proportionately, and that the heatwave exacerbates this scale effect while also introducing shape effects, causing a greater proportion of mosquitoes to reach older ages despite lowering the proportion of mosquitoes that survive the heatwave. Azacytidine, mostly eliminated these effects on the survival curves, and we argue that it is not because of the lower prevalence of infected mosquitoes. This is evident from our observations of DENV-infected mosquitoes treated with azacytidine (DENV2+A), which exhibited survival rates similar to those of non-infected mosquitoes from day 0 onward. If the 30% of the mosquitoes still infected (DENV2+A) had died at increasing rates as the infected mosquitoes (DENV2), it would have been reflected in the survival rates; however, this was not the case. We can say the same for the infected mosquitoes that were exposed to a heatwave and treated with azacytidine (DENV2+H+A vs. DENV2+H). At 7dpi, when the infection was evaluated, the difference in the survival rates between DENV2+H+A and DENV2+H was only 5%, which falls short if the 41% difference in the infection prevalence would be attributed to lower survival rates of the infected mosquitoes. Meanwhile, mosquitoes exposed to a heatwave have higher infection prevalence, which in combination with a longer life expectancy represents an increased potential risk for DENV transmission.

### 3.4 Heatwave increases the percentage of 5mC in total RNA in the *A. aegypti* midgut despite treatment with azacytidine

To determine whether the observed effects on survival rates and infection prevalence are related to the percentage of 5mC in RNA, we quantified the 5mC content in total RNA extracts by HPLC-FLD using 2-bromoacetophenone as a derivatizing agent. This agent fluorescently labels cytosines regardless of their methylation status. The 5mC percentage of total cytosines was measured at three time points: (1) 1 dpi in the midgut, time at which DENV infection is established and begins to spread within the midgut; (2) 4 dpi in the midgut, time at which DENV overcomes the midgut barrier and begins to spread to other organs; (3) 7 dpi in the head, thorax and abdomen, time at which DENV has successfully spread to other organs and body regions.

As can be seen in Figure 2A, the percentage of 5mC is 4.7 times higher in the midgut of mosquitoes exposed to a heatwave for one day (Mock 0.67% vs H 3.2%; p<0.0001), even when treated with azacytidine (H 3.2% vs. H+A 2.69%; ns). For its part, DENV2 increased the percentage of 5mC non-significantly (Mock 0.67% vs DENV2 0.98%; ns).

**Figure 2.**
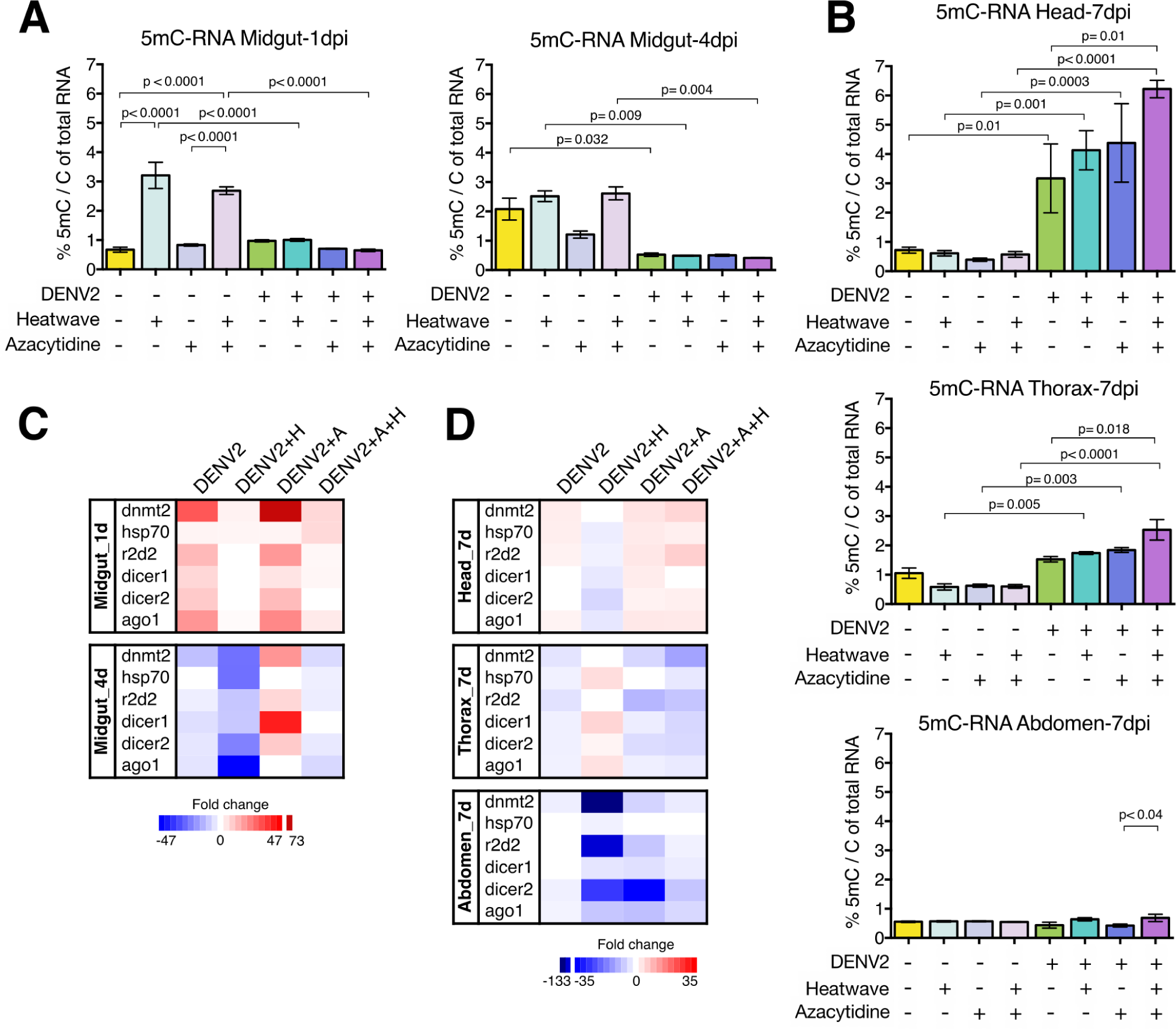
5mC content and transcriptional profile in *A. aegypti* challenged with DENV2 and exposed to heatwave or azacytidine treatment. 5mC content in total RNA from *A. aegypti* challenged with DENV2 and exposed to heatwave or azacytidine treatment. Total RNA was extracted from 10 specimens per condition and time. 2 µg of total RNA was derivatized with 2-bromoacetophenone and fluorometrically detected at 306/378 nm excitation/emission by HPLC-FLD. The percentage of 5mC was calculated relative to total cytidines in RNA. **(A)** 5mC content in total RNA from *A. aegypti*midgut at 1 dpi and 4 dpi. Representation of the arithmetic mean ± SEM of the overall 5mC percentage of three to five biological replicates performed in duplicates. ANOVA and Tukey’s multiple comparison test. Only *p-values*< 0.05 are shown. **(B)** 5mC content in total RNA from abdomen, thorax and head of *A. aegypti* 7 dpi. Plot of the arithmetic mean ± SEM of the overall 5mC percentage of three to five biological replicates performed in duplicates. ANOVA and Tukey’s multiple comparison test. Only *p-values*< 0.05 are shown. Transcriptional response of *A. aegypti* challenged with DENV2 and exposed to heatwave or azacytidine treatment. Heat map of transcriptional expression of *dnmt2*, *hsp70*, *r2d2*, *dicer1*, *dicer2* and *ago1* by qPCR. The color gradient represents the ratio (fold change) between the expression of the treated and control condition. Only genes with a fold change ≥ 2 or ≤ -2 were included in the heat maps; genes not meeting this criterion are shown in white boxes. **(C)** Relative expression of *dnmt2*, *hsp70* and antiviral iRNA pathway genes in midguts of *A. aegypti* 1 dpi and 4 dpi by qPCR. Representation of the mean of three biological replicates of 30 midguts each and two technical replicates. **(D)** Relative expression of *dnmt2*, *hsp70* and antiviral iRNA pathway genes in abdomen, thorax and head of *A. aegypti* 7 dpi by qPCR. Representation of the mean of three biological replicates of 30 specimens each and two technical replicates.

Stressful and nutritional stimuli increase 5mC content in RNA, with heatwave and blood feeding groups. The 5mC content in midguts of uninfected mosquitoes (Mock) increases 3-fold, from 0.67% (1dpi) to 2.08% (4 dpi), reflecting the physiological dynamics of 5mC in the gut after blood feeding (Figure 2A, 1 and 4 dpi).

### 3.5 DENV2 reduces the percentage of 5mC in total RNA in mosquito midguts, even if they were exposed to a heatwave

Oral exposure with DENV2 decreases 3.95-fold the percentage of 5mC in total RNA in 4 dpi midguts (Mock 2.08% vs. DENV2 0.53%; p = 0.032) and counteracts the heatwave-induced increase at 1 dpi (H 3.2% vs. DENV2+H 1.01%; p < 0.0001) (H+A 2.69% vs. DENV2+H+A 0.65%; p < 0.0001) and 4 dpi (H 2.52% vs. DENV2+H 0.49%; p = 0.009) (H+A 2.61% vs. DENV2+H+A 0.42%; p = 0.004). Thus, the percentage of 5mC in RNA remains low and constant in all experimental conditions where the mosquitoes were orally exposed to DENV2.

### 3.6 DENV increases the percentage of 5mC in total RNA in the mosquito head and thorax

The effect of DENV2 in the mosquito head and thorax is opposite to that observed in the midgut. While DENV counteracts the increase in the percentage of 5mC in the midgut (heatwave and azacytidine), it increases it more than 4-fold in the head and 3-fold in the thorax 7 dpi, especially with respect to the group of mosquitoes exposed to a heatwave and azacytidine (DENV2+H+A; Figure 2B).

At 7 dpi, the percentage of 5mC is less than 0.7% in mosquitoes exposed to a heatwave (H), treated with azacytidine (A) or that received both interventions (H+A), and is constant between abdomen, thorax and head (Figure 2B), meaning that there were no lasting effects on the percentage of 5mC 48 h after the interventions ended. The exception being the infected mosquitoes that were also exposed to a heatwave and treated with azacytidine (DENV2+H+A), in which there was a significant increase in 5mC content in head (DENV2 3.18% vs. DENV2+H+A 6.22%; p = 0.01) and thorax (DENV2 1.53% vs. DENV2+H+A 2.53%; p = 0.018) even though these interventions ended 48 h earlier.

The variation in %5mC at 7 dpi was segment dependent; in the abdomen the 5mC content ranged between 0.4 - 0.7%, in the thorax between 0.6 - 2.5% and in the head between 0.4 - 6.3% The abdomen was the segment with the lowest %5mC in all experimental conditions, while %5mC in thorax falls between that of the head and abdomen. The data in Figure 2B suggest that 5mC dynamics is positively related to the dynamic of DENV2 spread through the mosquito, beginning in the abdomen and progressing toward the head. As a potential proviral factor, 5mC levels temporarily increase during the initial phase of infection, returning to basal levels afterwards, presumably as host cells adapt to the virus.

### 3.7 The antiviral system is under-expressed in infected mosquitoes exposed to a heatwave

Next, we assessed the transcriptional profile of key members of the iRNA pathway as well as *dnmt2* and *hsp70* by RT-qPCR in the midgut of infected mosquitoes 1 dpi and 4 dpi compared to uninfected mosquitoes (Mock). As shown in the heat map (Figure 2C), the relative expression of the transcripts evaluated in midguts increases at 1 dpi when mosquitoes are orally challenged with DENV2 (fold change of +31 for *dnmt2*; +3 *hsp70*; +14 *r2d2*; +8 *dicer1*; +11 *dicer2*; +20 *ago1*), and is down-regulated 4 days after challenge (-11 *dnmt2*; -2 *r2d2*; -5 *dicer1*; -4 *dicer2*; -4 *ago1*), except *hsp70* which did not undergo any change. The under-expression of iRNA pathway members makes sense when considering the high prevalence of infection that *A. aegypti* develops.

Infected mosquitoes exposed to a heatwave (DENV2+H) exhibited a marked down-regulated transcriptional profile. At 1 dpi, marginal changes in the abundance of *dnmt2* (+3), *hsp70* (+3) and *ago1* (+2) were observed, whereas at 4 dpi, all transcripts were under-expressed (*dnmt2* -26; *hsp70* - 26*; r2d2* -10; *dicer1* -9; *dicer2* -23; *ago1* -47). These results indicate that exposure to a heatwave profoundly alters the gut transcriptional profile, down regulating the iRNA pathway in mosquitoes that develop 78% prevalence of viral infection. Notably, the use of azacytidine (DENV2+A) increases overall expression levels at 1 dpi (*dnmt2* +73; *hsp*70 +3; *r2d2* +20; *dicer1* +6; *dicer2* +13; *ago1* +23), although the pattern in wich expression levels decrease at 4 dpi (*dnmt2* +20; *r2d2* +8; *dicer2* +11) is maintained, except for *dicer1* (+42) which has a higher abundance at 4 dpi. Thus, the expression profile in the midgut of *A. aegypti* induced by DENV2 increases when mosquitoes are treated with azacytidine, transforming from a down-regulated to up-regulated transcript profile. The overexpression of members of the iRNA pathway is consistent with the low prevalence of infection in the group treated with the methylation inhibitor. Finally, the antagonistic effects on the expression profiles of azacytidine (DENV2+A) and heatwave (DENV2+H) appear to counteract each other when combined (DENV2+H+A), retaining the pattern in which an early response is induced (1 dpi) although marginal, and at 4 dpi is down-regulated, with the exception of *dicer1*, which did not change.

We then analyzed the transcriptional profile of the antiviral pathway, *dnmt2* and *hsp70* in the abdomen, thorax and head of mosquitoes 7 dpi, the time in which the virus had already spread to extra-abdominal regions compared to mock-infected mosquitoes. The relative expression levels are particular to body segment, with the abdomen exhibiting the lowest and the head the highest expression levels. DENV2 infection induces an upregulated transcriptional profile in the head (+2 to +3), whereas in the thorax (-2 to -5) and abdomen (-2 to -3) it is downregulated. These results are consistent with the dynamics of the transcriptional profile in the midgut, where there is a transcriptional overexpression in the organ at the time corresponding to the establishment of infection, and a transcriptional under expression at the time corresponding to viral dissemination to other organs. Consistent with the distinctive heat wave-induced transcriptional profile in gut, the most notable changes in transcriptional profiles in abdomen, thorax and head are observed in heat wave-exposed infected mosquitoes (DENV2+H), which contrast with the transcriptional profiles of the rest of the experimental groups. In the abdomen, heat wave reduced the expression of *dnmt2* (-133; fold), *r2d2* (-45) and *dicer2* (-27), which was also observed in azacytidine-treated mosquitoes (DENV2+A: *dnmt2* -8; *r2d2* -10; *dicer2* -35). In the abdomen, heatwave reduced the expression of *dnmt2* (-133; fold), *r2d2* (-45) and *dicer2* (-27), which was also observed in azacytidine-treated mosquitoes (DENV2+A: *dnmt2* -8; *r2d2* -10; *dicer2* -35). In the thorax, transcriptional overexpression occurred in mosquitoes exposed to a heatwave (DENV2+H: +2 to +7), contrasting with the down-regulation observed in all other experimental conditions. In contrast, expression changes in the head of mosquitoes exposed to a heatwave (DENV2+H) were down-regulated (-2 to - 7), a profile again divergent from the up-regulated patterns of the other experimental conditions.

Exposure to a heatwave (DENV2+H) and treatment with azacytidine (DENV2+A) caused profound and long-lasting changes in the transcriptional profile of mosquitoes as observed in the abdomen, thorax, and head two days after the end of the interventions. Exposure to a heatwave has the most notable effects on *dicer2* (abdomen -27; thorax +2; head -7), *r2d2* (abdomen -45; head -2) and *dnmt2* (abdomen -133). In infected mosquitoes exposed to a heatwave treated with azacytidine (DENV2+H+A) the most extensive changes in the transcriptional profile are observed in the thorax (-4 to -16; fold) and head (up-regulated +3 to +9) and share the most notable changes on *dicer2* (abdomen -10; thorax -7; head +3), *r2d2* (abdomen -2; thorax -10; head +9) and *dnmt2* (abdomen -3; thorax -16; head +7). The most noticeable effects of azacytidine treatment are observed on *dicer2* (abdomen -35; thorax -6; head +4), *r2d*2 (abdomen -10; thorax -13; head +4) and *dnmt2* (abdomen - 8; thorax -7; head +5).

### 3.8 Co-expression analyses reveal *dnmt2*, *r2d2*, and *ago1* as the most correlated genes

We constructed un-weighted co-expression analysis networks for each experimental condition and body segment combination to explore possible relationships in the differential relative expression of the genes of interest across various tissues and experimental conditions (44). The relative expression against the control group was used to compute the Pearson correlation matrices for the genes in each tissue and experimental condition, which were then filtered by the *p-*value of the correlation with a threshold cut-off of *p ≤ 0.05*. The networks were more complex in the experimental conditions that included azacytidine (DENV2+H, DENV2+A, DENV2+H+A) concerning to the infected mosquitoes alone (DENV2) (Figure 3A). Although the infected mosquitoes that were treated with azacytidine and exposed to a heatwave (DENV2+H+A) showed the highest number of represented genes or nodes, the correlations were less than in the infected mosquitoes solely treated with azacytidine (DENV2+A), which presented 16 nodes and 17 edges (Figure 3A). This means that there were less co-expression relationships in DENV2+H+A mosquitoes. On the other side of the spectrum are the infected mosquitoes (DENV2), which showed 11 nodes with 7 edges, and only one network comprised of more than 2 nodes in the head. The most complex networks were observed in the infected mosquitoes treated with azacytidine (DENV2+A, DENV2+H+A), especially in the 1 dpi midgut and 7 dpi thorax.

**Figure 3.**
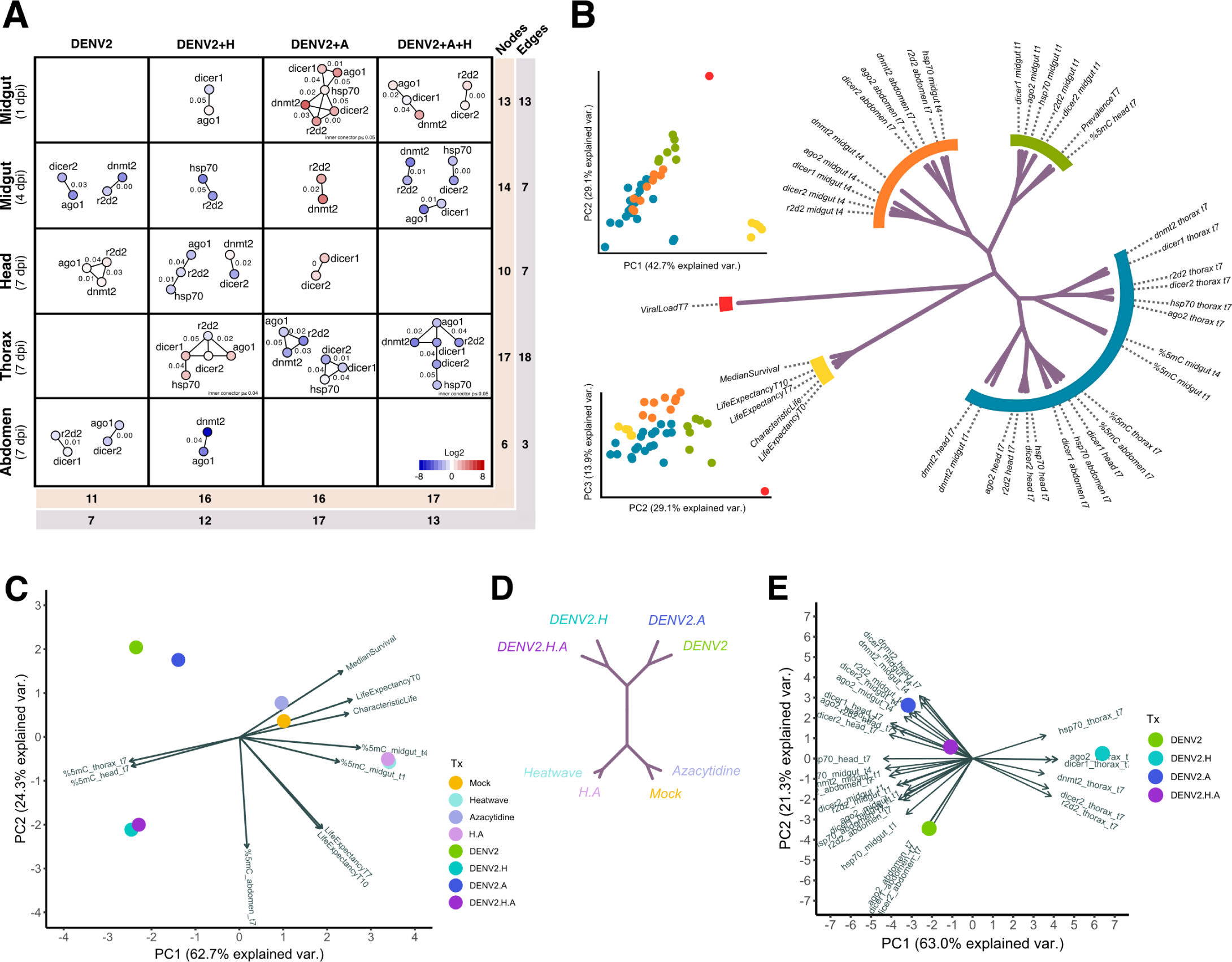
Co-expression and principal component analysis. **(A)** Transcriptional co-expression analysis of *dnmt2*, *hsp70*, *r2d2*, *dicer1*, *dicer2*, and *ago1* in *A. aegypti* mosquitoes challenged with DENV2 and exposed to a heatwave or treated with azacytidine. RT-qPCR data were normalized with respect to *RNase P* expression (2-Δ^Ct^), Log2-transformed, and relativized to the control group before calculating the Pearson correlation matrix. Significant correlations were filtered using a cutoff threshold of p ≤ 0.05, which resulted in correlations with R2 ≥ 0.996 or ≤ -0.995. Blank boxes indicate that no significant correlation was found. The number of nodes (genes) and edges (correlations) per body segment or experimental condition are shown at the sides and bottom. The color scale represents relative expression levels (Log2). The *p* value is shown as numbers adjacent to the interactions. **(B)** Principal component analysis. Data from all biological replicates generated in this study were averaged prior to analysis. Clusters in the dendrogram were generated by hierarchical analysis from the Euclidean distance between the coordinates of the observations in the principal component space. The resulting tree was cut into 5 branches to define the biggest clusters. **(C)** PCA of experimental conditions. Data were transposed to show relationships between experimental conditions. Viral load and infection prevalence variables were omitted to avoid introducing bias during clustering. Arrows indicate the variables, how they correlate with each other and how they are described by the principal components. Variables are correlated if they are collinear and uncorrelated if they are orthogonal to each other. The vectors that make up each variable describe how it contributes to the principal components. **(D)** Treatment clustering. The variables analyzed in (C) allowed differential clustering of infected mosquitoes from uninfected mosquitoes. heatwave-exposed mosquitoes subclustered within the infectious status clusters. **(E)** PCA of the relative expression data. DENV2+H mosquitoes form their own cluster while DENV2+A, and DENV2+H+A form another cluster, indicating that the azacytidine treatment exerted greater influence on gene transcription than the heatwave alone. The contributions in gene expression of azacytidine and heatwave treated mosquitoes antagonized each other (DENV2+H+A lies close to 0 in both x and y axis, indicating very low variation in the data compared to the control group to which it is relativized.

Regarding the body segments and midgut, the thorax showed the most complex networks in terms of both the number of nodes and edges, while the abdomen presented the least. In addition, the 4 dpi midgut and head showed a disparity between the number of nodes and edges. This was because the co-expression networks were less intertwined, forming many small separate networks of 2 or 3 nodes. It stands out from the comparisons that the body segments/midgut tissue lack significant co-expression relationships in at least one of the combinations (empty boxes in Figure 3A), with the exception being 4 dpi midguts. Of particular interest is the abdomen, where almost no co-expression was seen. In contrast, heatwave-exposed infected mosquitoes (DENV2+H) presented co-expressed genes in all body segments and midgut.

From the analysis of individual genes and interactions, it stands out that in 4 dpi midguts, *dnmt2* correlated with *r2d2* in all experimental conditions except in the heatwave-exposed mosquitoes (DENV2+H), where instead *r2d2* co-expressed with *hsp70*. Similarly, the co-expression of *r2d2* and *ago1* predominated in the head and thorax (Figure 3A). Interestingly, *dnmt2* and *r2d2* were the two most co-expressed genes across the samples with 6 different instances of co-expression (3 of them in the 4dpi midgut), just followed by *r2d2* and *ago1* with 5 instances of correlation, and *dicer1* and *ago1*, also with 5 (3 of them in DENV2+H+A mosquitoes). This makes it so the most co-expressed genes, regardless to which they are correlated to, are *r2d2* with 19 edges, *ago1* with 18 edges and *dnmt2* with 15 edges in total. *hsp70* showed only 12 instances of co-expression among the samples, suggesting that *hsp70* expression is fine tuned independently. In fact, *hsp70* was not found correlated neither in the infected (DENV2) nor in the abdomen in any experimental condition. Notable are the co-expression relationships between *hsp70* and *ago1*, as well as *hsp70* and *dnmt2*, which were exclusively observed in the 1 dpi midgut of DENV-infected and azacytidine-treated mosquitoes (DENV2+A), unlike the co-expression of *dnmt2* and *ago1*, that was seen under all experimental conditions except in the infected and azacytidine treated mosquitoes (DENV2+A) (Figure 3A).

Most of the co-expression relationships observed were positive correlations, where the co-expressed genes were either upregulated or downregulated in relation to the control (Figure 3A). The only few inverse correlations were found in 1 dpi midguts of infected mosquitoes treated with azacytidine (DENV2+A), and also in those additionally exposed to a heatwave (DENV2+H+A), where *dicer1* was downregulated while *ago1* was upregulated. In DENV2+H+A mosquitoes, *dnmt2* was also co-expressed with the latter genes. The head of infected and azacytidine-treated mosquitoes (DENV2+A) presented also another case of inverse correlation between *dnmt2* (upregulated) and *dicer2* (downregulated). Finally, inverse correlations were also found in the thorax of infected and azacytidine-treated mosquitoes as well, where *r2d2* (downregulated) was inversely correlated with *dicer1*, *dicer2*, and *ago1* (upregulated) (Figure 3A).

### 3.9 How mosquitoes respond to a heatwave depends on whether they are infected with DENV

To integrate and find response patterns in the data, we performed principal component analysis (PCA) to assess the contributions of the experimental conditions in explaining the differences in the evaluated factors. We then calculated the Euclidian distance between the factors and constructed a hierarchical dendrogram (Figure 3B). The factors clustered into 5 major distinct groups, which mean that the factors behaved in 5 different general ways. For example, the infection prevalence and the 5mC content in the head subclustered together within the green cluster because the infected mosquitoes showed high levels of 5mC and prevalence, while uninfected mosquitoes showed low 5mC levels and prevalence; both variables have an inverse relationship with the uninfected condition and a positive relationship with the infected condition. The other genes in the green cluster showed similar but divergent response patterns. Similarly, in 1 dpi midguts, the expression pattern of *dnmt2* (blue cluster) showed greater similarity to its expression in the head, as well as to other genes expressed in the head, rather than its expression to other genes within the same body segment. However, this represents an exceptional case where a gene displayed consistent responses irrespective of its body segment, as the majority of genes showed expression patterns based on the body segment or tissue. Genes tended to cluster together on the basis of body segment, rather than on the basis of what gene they are.

We then asked how these factors contributed to explain the differences in the experimental conditions. This would allow us to find shared response patterns between the experimental conditions instead of the factors’ particular responses. For this, we performed another PCA but now considering the experimental conditions as factors (Figs. 3C and 3E). Since the contribution of the viral load and prevalence in explaining the differences between the experimental conditions is straightforward, we ditched them for further analysis. This would also allow us to determine patterns between infected and non-infected mosquitoes beyond considering their infection status. We also analyzed the gene expression data separately as the relative expression because it presented its own particular behavior. As seen in Figure 3C, without considering the infection parameters and gene expression patterns, DENV2 induced a distinct overall response in the mosquitoes that distinguished them from the non-infected mosquitoes, separating the experimental conditions in two clusters, an infected one and an uninfected one (Figure 3D). The heatwave also induced a characteristic, albeit weaker response in the mosquitoes forming two smaller clusters, one within the infected group and the other one within the uninfected group. The azacytidine treatment induced a heterogeneous response that depended on whether the mosquitoes were infected or exposed to a heatwave; therefore, these experimental groups lay within separate clusters. Life expectancy and the %5mC in abdomens at 7 dpi explain the differences in the second principal component of the infected mosquitoes, while the median survival, characteristic life, and %5mC in other segments/midgut explain the uninfected condition in the first principal component. Finally, the PCA of the relative expression bears interest in that the infected mosquitoes that were exposed to a heatwave, behaved transcriptionally in an opposite manner as the other experimental conditions, thus contributing the most to the variation in the data (Figure 3E). While most of the genes in the different body segments presented some degree of positive association with each other, the transcriptional response in the thorax was the opposite, presenting negative correlation with the rest of the variables as denoted by the obtuse angles between the thorax and other tissues or body segments (arrows in Figure 3E).

In conclusion, we have determined how the mosquitoes respond to a DENV infection in a scenario where a heatwave is also happening, and how the disruption of 5mC influences this response. Azacytidine has profound and pleiotropic effects on mosquito physiology that lead to lower infection rates. We confirmed the significant impact DENV has on the mosquito and how changes in ambient temperature modify the mosquito response, ranging from global changes in the transcriptional profile, changes in the dynamics of 5mC, but above all, in the life expectancy, which represents a risk for the potential transmission of DENV. The final contribution of this work is the consideration of 5-methylcytosine as an additional factor in vector competence and DNMT2 as a potential target to disrupt DENV transmission.

## 4 Discussion

5-methylcytosine in RNA plays a pivotal role in regulating transcriptional expression, influencing a wide range of cellular processes, including energy and lipid metabolism, and responses to oxidative stress, heat stress, and the immune challenge (13,16,17,45–51). These processes are intimately linked to cellular adaptation to challenging environmental factors. While 5mC is found across the entire phylogeny, its role in biological processes in mosquitoes remains underexplored, particularly within the context of climate change.

Dipterans such as *Drosophila*, *Anopheles* and *Aedes* respond transcriptionally by increasing *dnmt2* methyltransferase expression to biotic (such as challenge with RNA viruses, bacteria and parasites) and abiotic (such as heat and oxidative stress) challenges (18–23,30,32,33,35). Consistent with the above, our results show that *dnmt2* expression increases in *A. aegypti* in response to DENV2 infection, heat stress (heatwave) and methylation inhibition treatment (azacytidine) (Figure 1A). Consistently, %5mC-RNA in midguts of DENV2-challenged *A. aegypti* mosquitoes increases to nearly 2-fold, is 3-fold higher when exposed to a heatwave, and remains at low levels when mosquitoes are treated with azacytidine despite *dnmt2* overexpression (+73-fold) (Figure 2A and 2C). These findings contribute to the growing body of evidence supporting *dnmt2* as a universal factor involved in the stress response in mosquitoes, and they underscore the elevation in 5mC-RNA content as a response to challenging environmental factors.

Previous studies have suggested that *dnmt2* is a proviral factor in mosquitoes. Research conducted by Zhang (18) and Bhattacharya (22) delves into the multifaceted roles of *dnmt2*, addressing its interactions within intricate scenarios involving insects, pathogens and symbionts. Zhang’s work reveals that DENV infection induces *dnmt2* expression, whereas *Wolbachia* suppresses it by expressing the microRNA aae-miR-2940. Notably, overexpression of *dnmt2* in mosquito cells inhibits *Wolbachia* growth but enhances DENV replication. Bhattacharya et al. (2022) also reported that *dnmt2* exhibits proviral function in alpha viruses such as Sindbis (SINV) and Chikungunya virus (CHIKV), as it increases their infectivity in cells overexpressing DNMT2. In agreement, our results support the hypothesis that *dnmt2* is a proviral factor in mosquitoes. We found that *dnmt2* expression increases and 5mC-RNA tends to increase (Mock 0.67% vs. DENV2 0.98%; ns) in the midgut of *A. aegypti* mosquitoes at 1 dpi, when viral replication is active in the midgut(40). As the virus disseminates from the midgut to other organs at 4 dpi, *dnmt2* expression and 5mC-RNA levels decrease, and by 7 dpi, the expression of *dnmt2* is already downregulated and 5mC returns to basal levels in the abdomen. In the thorax, *dnmt2* is also downregulated and the 5mC returns to basal levels, while the head seems to be the site of active replication showing *dnmt2* upregulation and a 4-fold increase in 5mC levels. These results suggest that RNA methylation in *A. aegypti* acts as a proviral factor that could be related to the regulation of viral replication. DNMT2 may methylate host RNA, which could alter gene expression and create a more favorable environment for viral replication. Using LC-MS, Bhattacharya et al. (2022) (22)identified 5mC in SINV and CHIKV RNA obtained from mosquito cells (C7/10, 5 dpi), and proposed that these alpha viruses are targets of DNMT2 methylation. Although quantification by HPLC-FLD has a detection limit on the order of femtomoles, we did not find 5mC in viral RNA from infected C6/36 cells at 7 dpi (Supplementary Figure 1B), indicating that the 5mC measured in our samples was derived from the mosquito itself.

In this work we opted for a pharmacological inhibition approach using azacytidine to target RNA methylation, specifically inhibiting C5 methyltransferases (C5-MT; Reviewed in: (26,52). In the presence of inhibitors, RNA C5-methyltransferases (C5-RMT) are overexpressed, and eventually the amount of 5mC in RNA decreases (53). In our study, treatment with the RNA methylation inhibitor led to a substantial increase in *dnmt2* expression, low 5mC content 4 days after treatment and clear biological effects on transcriptional profile. Both azacytidine methylation inhibitor treatment and heat stress produce profound and complex changes in the cellular transcription profile (53,54). We found that the expression profile in the midgut of infected mosquitoes is lower if exposed to a heatwave and higher if treated with azacytidine. Moreover, these trends in transcript expression persist in the head two days after the stimuli ceased. Thus, infected mosquitoes exposed to a heatwave downregulate the transcriptional profile, while those treated with azacytidine upregulate it.

Stress responses are triggered by the impairment of protein function due to stressors such as temperature fluctuations, UV-radiation, infections, and oxidative stress, resulting in notable changes in the overall transcriptional profile. In the case of heat stress, alterations in the transcriptional profile drive the synthesis of Hsp chaperones, which play a critical role in preventing protein misfolding and polypeptide aggregation. However, the production of Hsp chaperones comes with a cost, as their accumulation can lead to toxicity, indicating that the relationship between Hsp production and the stress response can be multifaceted. In *A. aegypti* mosquitoes, the heat stress response, characterized by moderate *hsp* expression, is associated with enhanced thermal tolerance and effective adaptation to heat stress (55). In this context, Ware-Gilmore et al. (2023) proposes that insect responses characterized by high or sustained chaperone production may indicate an inadequate adaptation to heat stress.

The expression of the multifunctional protein Hsp70 plays a pivotal role in various cellular processes. In *A. aegypti*, Hsp70 is required for DENV cellular entry, RNA replication and virion biogenesis, thereby earning its status as a vector factor with proviral functionality (56). Indeed, oral challenge with DENV2 leads to an increase in *hsp70* expression, displaying similar dynamics to those observed in *dnmt2*, in which gene transcription in the midgut, abdomen, thorax, and head may be associated with active DENV2 replication. While the relative expression of *hsp70* in the midgut exhibited a 3-fold increase following DENV infection at 1 dpi, no additional effect was observed upon exposure to a heatwave. Relative *hsp70* expression did not exceed a 3-fold change in any segment/tissue 7 dpi and after a heatwave exposure (DENV2+H), the *hsp*70 response was moderate, with changes of -3 (head), +6 (thorax) and -2 (abdomen) fold. The exception were the 4 dpi midguts, in which *hsp70* expression decreased considerably (-26-fold), coinciding with the period of highest mortality of mosquitoes exposed to the heatwave (DENV2+H; Figure 1C). These results indicate that *A. aegypti* does not respond disproportionately with *hsp70* to the heatwave and that the decrease in *hsp70* could be related to the higher mortality observed.

Variables such as the virus strain, viral load and route of infection play crucial roles in determining the infection success and its impact on the survival (fitness) of infected mosquitoes, particularly in challenging environmental factors. Our results show that the DENV2 infection, exposure to a heatwave, and azacytidine treatment significantly affect *A. aegypti* survival. Firstly, DENV2 infection, lead to a substantial reduction in mosquito survival, a trend consistent with previous studies highlighting the detrimental impact of viral infection on mosquito survival (57). Interestingly, the adverse effects of DENV2 infection on survival are reversed when mosquitoes are treated with azacytidine. This finding suggests that azacytidine antagonizes *dnmt*2 as a proviral factor, favoring the ability of mosquitoes to resist infection. Future research should explore the modulation of the immune response or the reduction of viral replication as potential mechanisms underlying these effects. Secondly, we observed a reduction in mosquito survival when infected with DENV2 and exposed to a heatwave. This observation aligns with previous studies, such as the work by Maciel-de-

Freitas et al. (2010) (57), which demonstrate that elevated temperatures can exert a negative impact on the survival and reproductive capabilities of *A. aegypti* mosquitoes. This synergistic effect between viral infection and heat stress holds significance in understanding how environmental conditions can affect dengue virus transmission. Thirdly, we observed that even with azacytidine treatment, the survival of DENV2-infected mosquitoes exposed to a heatwave remained significantly diminished. This finding highlights that azacytidine is not able to fully compensate for the negative effects of the infection on the survival of mosquitoes exposed to a heatwave. Conversely, mosquitoes unchallenged with DENV2 but subjected to azacytidine treatment, heatwave exposure, or the combination of azacytidine and heatwave, exhibited no difference in survivorship from the control group by the Log-rank test. These results align partially with prior research, such as that of Christofferson et al. (2016) (58), in which varying temperatures (26°C, 28°C and 30°C) had no impact on the survival of uninfected mosquitoes, while in DENV2-infected mosquitoes survival is reduced only when held at the highest temperature (30°C). Complementarily, we analyzed the effects of DENV2 infection, exposure to a heatwave, and azacytidine treatment on the life expectancy of *A. aegypti* mosquitoes. Consistent with the survival rates, the estimated life expectancy is lower for DENV2-infected mosquitoes, indicating that viral infection exerts a detrimental effect on mosquito biology. The most striking observation is that DENV2-infected mosquitoes showed an increased life expectancy when exposed to a heatwave. This result prompts inquiries into the thermal adaptation mechanisms employed by mosquitoes and how these adaptations might influence their potential for virus transmission. Since temperature accelerates the extrinsic incubation period of DENV (9). infected mosquitoes exposed to a heatwave, in which a longer life expectancy is observed, may engage in more feeding episodes, thereby introducing an elevated risk associated with DENV transmission.

Recent research by Wimalasiri-Yapa et al. (2021), demonstrated that infected mosquitoes maintained at 32°C, the highest temperature tested, exhibit elevated virus loads and a diminished transcriptional response in immune pathway genes (59). Correspondingly, we have shown that heatwave-exposed mosquitoes display a suppressed transcriptional profile in antiviral pathway genes and develop high infection rates. Interestingly, the life expectancy of infected mosquitoes exposed to a heatwave is reduced when concurrently treated with the methylation inhibitor. Methylation of tRNA is known to be inducible, with *dnmt2* playing a pivotal role in mediating an adaptive response, particularly under conditions of heightened metabolic demand, such as during stress periods(33,60). Transcriptional responses in DENV-infected mosquitoes are enriched in transcripts associated with metabolic processes, redox functions, and ATP generation (39,61). 5mC is also recognized for its involvement in various metabolic processes, including protein synthesis, energy metabolism, and lipid metabolism (15–17,46–49,51). The 5mC at C38 in tRNA-Asp is known to be related to three critical mechanisms: (1) an increased rate of tRNA/amino acid loading by aspatyl tRNA synthetase, (2) enhanced tRNA stability, and (3) improved fidelity in anticodon/codon interaction, minimizing mistranslation of glutamate to aspartate codons (as reviewed in: (28). Thus, the role of methylation in the context of inhibition may compromise protein synthesis.

Finally, the prevalence of DENV2 infection in mosquitoes is reduced by 53% when treated with azacytidine, indicating a potential role for methylation in the viral infection. *A. aegypti* mosquitoes exposed to a heatwave showed high viral loads and DENV2 prevalence, but when subjected to azacytidine treatment, the infection prevalence decreased by 41%. This finding suggests that azacytidine may mitigate the negative effects of heat exposure on virus transmission. This notion finds support in the work of Gale (2019)(62), whose study showed that elevated temperatures increases vector competence of *A. aegypti* mosquitoes for DENV. Temperature affects various aspects of the virus-mosquito interaction, including the binding affinity of the virus to host cells, the rate of viral replication, and its spread to the salivary glands.

In summary, our results strongly suggest that *dnmt2* and RNA methylation play integral roles in the response of *A. aegypti* to both heat stress and dengue virus infection. RNA methylation appears to play a complex role in mosquito biology and in their interaction with the virus. Our findings demonstrate that inhibition of RNA methylation with azacytidine reduces the infection prevalence and improves mosquito survival under heat stress conditions, which may have far-reaching implications for dengue transmission dynamics. Nevertheless, further investigations are imperative to comprehensively unravel the underlying mechanisms and assess the full potential of this strategy for controlling mosquito-borne diseases.

## 6 Author Contributions

FCP conceptualized the project, designed and performed the experiments, analysed and interpreted the data, and wrote the original draft. BRT performed the experiments, analysed and interpreted the data, and wrote the original draft. HLM and JCC administrated the project, provided resources, supervised the project, and wrote, revised, and edited the draft.

## 8 Acknowledgments

This is a short text to acknowledge the contributions of specific colleagues, institutions, or agencies that aided the efforts of the authors.

